# Apnoea suppresses brain activity in infants

**DOI:** 10.1101/2024.02.16.580547

**Authors:** Coen S. Zandvoort, Anneleen Dereymaeker, Luke Baxter, Katrien Jansen, Gunnar Naulaers, Maarten de Vos, Caroline Hartley

**Affiliations:** Department of Paediatrics, University of Oxford, Oxford, United Kingdom; Department of Development and Regeneration, University Hospitals Leuven, Neonatal Intensive Care Unit, KU Leuven, Leuven, Belgium; Department of Development and Regeneration, University Hospitals Leuven, Child Neurology, KU Leuven, Leuven, Belgium; Department of Electrical Engineering (ESAT), STADIUS Center for Dynamical Systems, Signal Processing and Data Analytics, KU Leuven, Leuven, Belgium

**Author notes:** Corresponding author, Paediatric Neuroimaging Group, Department of Paediatrics, Level 2 Children’s Hospital, John Radcliffe Hospital, University of Oxford, Oxford, United Kingdom.

**Keywords:** *Neonate*, *brain*, *electroencephalography*, *respiration*, *early development*

## Abstract

Apnoea – the cessation of breathing – is commonly observed in premature infants. These events can reduce cerebral oxygenation and are associated with poorer neurodevelopmental outcomes. However, relatively little is known about how apnoea and shorter pauses in breathing impact brain function in infants, which will provide greater mechanistic understanding of how apnoea affects brain development. We analysed simultaneous recordings of respiration, electroencephalography (EEG), heart rate, and peripheral oxygen saturation in 124 recordings from 118 infants (post-menstrual age: 38.6 ± 2.7 weeks [mean ± standard deviation]) during apnoeas (pauses in breathing greater than 15 seconds) and shorter pauses in breathing between 5 and 15 seconds. EEG amplitude significantly decreased during both apnoeas and shorter pauses in breathing compared with normal breathing periods. Change in EEG amplitude was significantly associated with change in heart rate during apnoea and breathing pauses and, during apnoeas only, with oxygen saturation change. No associations were found between EEG amplitude and pause duration or post-menstrual age. The decrease in EEG amplitude may be a result of the changing metabolism and/or homeostasis following changes in oxygen and carbon dioxide concentrations, which alters the release of neurotransmitters. As apnoeas often occur in premature infants, frequent disruption to brain activity may impact neural development and result in long-term neurodevelopmental consequences.

## 1 Introduction

Worldwide, 15 million infants are born prematurely each year (WHO, 2023), putting them at increased risk of long-term neurodevelopmental disability. Apnoea frequently occurs in preterm infants, affecting more than half, and all infants born before 29 weeks’ gestation (Eichenwald et al., 1997; Finer et al., 2006). These events can result in cerebral deoxygenation (Choi et al., 2021; Horne et al., 2018) and recurrent episodes of apnoea have been associated with poorer long-term neurodevelopmental outcomes (Janvier et al., 2004; Pillekamp et al., 2007). Yet, it remains unclear the extent to which apnoea is causative or simply correlated with poorer later-life neurodevelopmental outcome (Williamson et al., 2021). Understanding the short-term impact of apnoea on the developing infant brain will shed light on this relationship. A comprehensive framework of the interaction between brain activity and apnoea is lacking but pivotal for gaining a better mechanistic understanding of how apnoea affects brain development in premature infants.

To date, a small number of studies have investigated the effects of apnoeic episodes on brain activity recorded using electroencephalography (EEG) in infants (Usman et al., 2023). Although the amplitude of brain activity decreases in some apnoeic episodes compared to periods of normal breathing (Bridgers et al., 1985; Deuel, 1973; Low et al., 2012; Wulbrand and Bentele, 1994), the overall findings of these studies are inconsistent (Usman et al., 2023). Limited sample sizes (i.e., <10 infants) have precluded investigation of factors which may modulate the effects and explain these inconsistencies (Usman et al., 2023). Whilst long apnoeic episodes can cause complete suppression of cortical activity and severe hypoxia (Low et al., 2012), duration may not be associated with the level of EEG suppression for shorter episodes (Fenichel et al., 1980). Moreover, it remains unclear how the infant’s age and levels of physiological instability (e.g., oxygen desaturation and bradycardia) come into play. In piglets, brain activity and oxygen saturation decreased following experimentally induced apnoea, but heart rate only decreased later (Sanocka et al., 1988). A similar pattern, with later occurring bradycardia compared with changes in EEG, was observed in a single human neonate recorded by Deuel (1973). Yet, the relationship between apnoea-related EEG changes and bradycardia and oxygen desaturation requires further investigation.

We aimed to study the acute effects of apnoea on brain activity in infants and its relationships with heart rate change, oxygen saturation change, breathing pause duration, and age. Following earlier reports, we hypothesised that EEG amplitude decreases during apnoea and shorter pauses in breathing. We build on these previous results by applying a quantitative approach to characterise the modulation of cortical activity during breathing pauses using time-frequency-amplitude analysis, and by examining how cortical activity changes relate to physiological instability, pause duration and age. Apnoea is often defined as cessation of breathing which lasts at least 15 seconds, or longer than 10 seconds when accompanied by bradycardia or cyanosis (Blackmon et al., 2003; Elder et al., 2013; Martin et al., 2004). We investigated how brain activity changes in response to apnoeic episodes of at least 15 seconds in duration. Moreover, since the apnoea definition of 15 seconds is somewhat arbitrary, isolated shorter pauses may elicit changes in neurophysiology and so we also investigated changes in brain activity following breathing pauses between 5 and 15 seconds. For clarity, we examined changes in brain activity during isolated pauses in breathing only. This is opposed to periodic breathing/clustered pauses where several short breathing pauses occur closely together.

## 2 Material and methods

### 2.1 Participants and study design

The dataset was acquired at the neonatal intensive care unit of the University Hospitals Leuven, Belgium. The study protocol was in accordance with guidelines on Good Clinical Practice and Declaration of Helsinki and approved by the ethics committee of the University Hospitals Leuven. Data were recorded for research purposes (Pillay et al., 2020; Pillay et al., 2018). Parents or legal guardians provided written and oral consent for their child’s data to be used for research purposes. Infants were included in the current study if they had both respiration and brain activity recordings. Data comprised a total of 124 recordings from 118 infants (112 babies had a single test occasion and six babies had two test occasions). Demographic details are provided in Table 1.

**Table 1.** Demographic details. Reported values are mean ± standard deviation [range]. Superscripts indicate number of days.

| **Factor** |  |
| --- | --- |
| Gestational age at birth (weeks) | $30.43 \pm 2.90$ [$24^{+4}$; $40^{+0}$] |
| Post-menstrual age at recording (weeks) | $38.56 \pm 2.66$ [$31^{+2}$; $47^{+5}$] |
| Chronological age at recording (weeks) | $8.13 \pm 3.94$ [$0^{+4}$; $22^{+1}$] |

### 2.2 Data acquisition

EEG was recorded at a sampling rate of 250 Hz using Brain RT OSG Equipment (Mechelen, Belgium). A 9-channel full-scalp configuration comprising Fp1, Fp2, C3, Cz, C4, T3, T4, O1, and O2 electrodes (positioned according to the international 10-20 system) was used to collect EEG data in 122 out of 124 recordings. In two recordings, subsets of four and five channels were used. Data were referenced to channel Cz. EEG data were recorded for an average of 15.63 ± 6.23 [1.31; 25.96] hours (median ± interquartile range [range]).

Respiration, heart rate and peripheral oxygen saturation were simultaneously acquired using Philips IntelliVue monitors. Respiration was recorded at the thorax (*n* = 124), abdomen (*n* = 114) and nasal airways (*n* = 103). All respiratory signals were sampled at 25 Hz. Heart rate and oxygen saturation signals were acquired at 1 or 5 Hz in 111 recordings (*n* = 47 recordings at 5 Hz and *n* = 64 at 1 Hz). Oxygen saturation was measured with a bandwidth of 60-100% with the sensor placed peripherally on the foot or hand.

### 2.3 Data analysis

All analyses were performed in MATLAB (ver. 2022b; MathWorks Inc., Natick, USA) and are summarised in Figure 1. In brief, we first identified apnoeas and shorter pauses in breathing from respiration signals. Next, we investigated changes in EEG amplitude during apnoea and shorter pauses in breathing compared with periods of normal breathing using time-frequency analysis. Finally, we compared the change in EEG amplitude with change in heart rate, change in oxygen saturation, duration of the breathing pause and age of the infant.

**Figure 1.**
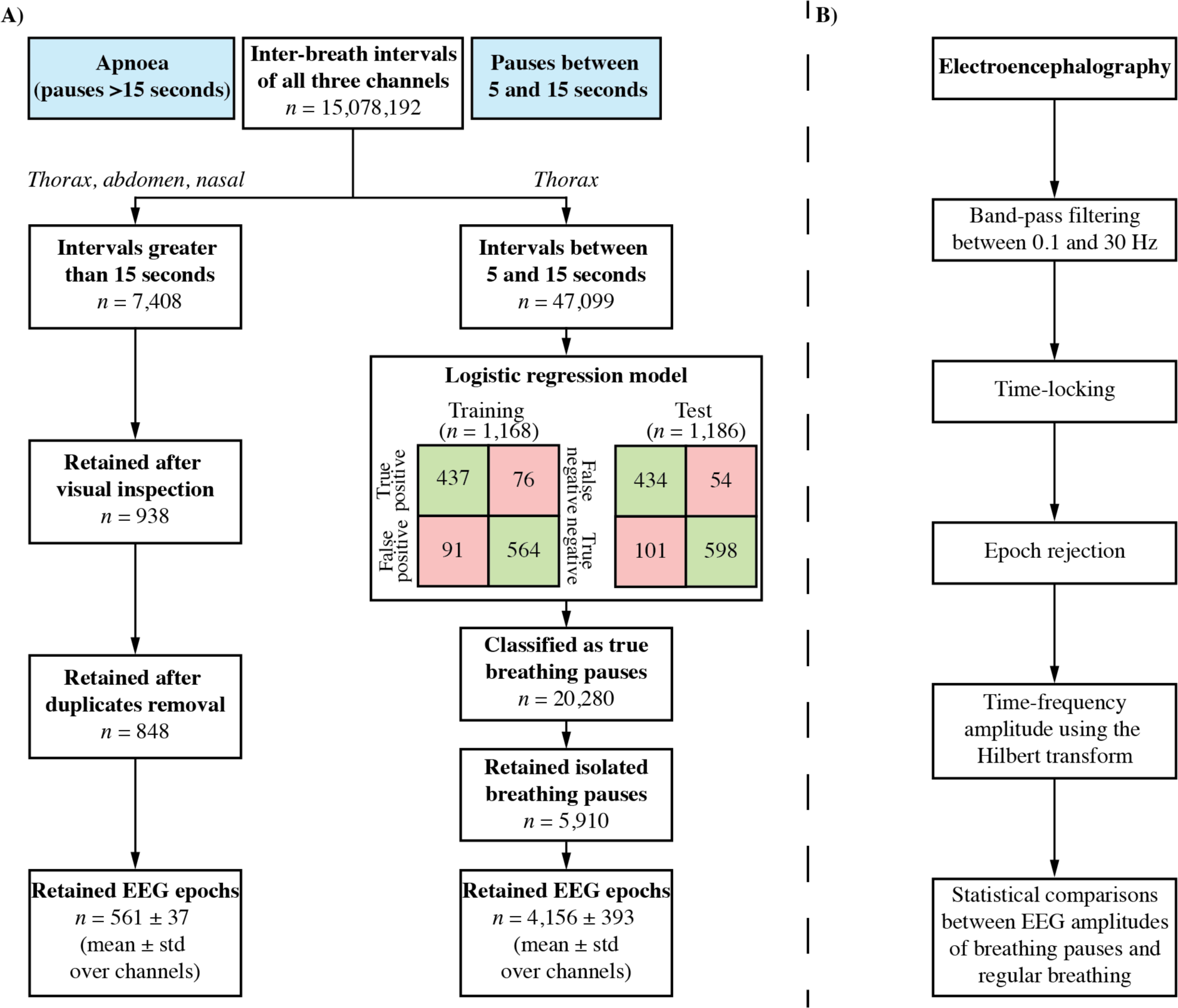
Flowchart of breathing pause identification and electroencephalographic analysis. A) Breakdown is provided for apnoeas (pauses greater than 15 seconds; left stream) and breathing pauses between 5 and 15 seconds (right stream). **B)** Analysis pipeline of the electroencephalography.

### 2.4 Breathing pause identification from the respiration signals

Figure 1A provides an overview of the steps taken for the identification of apnoea and shorter pauses in breathing. Respiratory signals were first high-pass filtered with a cut-off frequency of 0.5 Hz (bidirectional second-order Butterworth filter). After that, signals were checked for amplitude outliers. Samples that exceeded five times the standard deviation of the signal were removed and linearly interpolated, a process that was iterated 10 times. We then identified the inter-breath intervals (IBI) in each of the three respiratory signals separately using a previously described algorithm validated for the identification of IBI in infants from the impedance pneumograph (Adjei et al., 2021), adapting this algorithm for the respiratory signals recorded here. Briefly, the algorithm first removes artefacts from the respiration signals that could falsely be detected as breathing cycles. It then computes a threshold according to 0.4 times the signal’s standard deviation across the previous 15 breaths. Visual validation of the algorithm on the alternative respiratory signals used here led us to modify these parameter values to a threshold of 0.5 times the standard deviation over 120 breaths. Individual breaths are detected as times when the respiratory signal crosses this threshold. Finally, the published algorithm removes false positive apnoeas using a machine learning classifier. Due to the different signals used here we instead visually inspected all possible apnoeas (IBI>15 seconds) to discard periods of low amplitude signal falsely detected as apnoeas, for example, due to hypopnea/shallow breathing or artefact.

For breathing pauses between 5 and 15 seconds, we focused on the IBI identified from the thoracic signal, which had a higher signal-to-noise ratio than the abdomen and nasal airflow signals. From this signal, we identified 47,099 potential breathing pauses. As manually looking through each possible breathing pause is too time demanding, we developed a classification model that accurately identifies true pauses in breathing. Pauses in breathing between 5 and 15 seconds that were identified by the classifier as a true pause in breathing were included in the EEG analysis. To derive the classification model, 1,168 possible pauses were used as a training set and 1,186 possible pauses were used as a test set. An experienced researcher (CZ) labelled each of the possible pauses as true or incorrectly identified. We next computed the mean absolute and standard deviation of the thoracic signals in three time windows. Time windows consisted of the 10 to 1 seconds before and 1 to 10 seconds after the pause in breathing. The third window was from 1 second after the start of the pause in breathing to 1 second before the end of the pause in breathing. To account for amplitude differences across infants, infant-specific thoracic signals were normalised to the standard deviation of the full recording. Using the researcher’s ratings, a logistic regression model was trained yielding a balanced accuracy of 86% for the training set. Applying this model to the test set gave a balanced accuracy of 87% demonstrating the applicability of the model to independent data. From all 47,099 potential pauses, the logistic regression classified 20,280 of them as true. To rule out any effect of short pauses in breathing in the baseline or post-pause period, we focused our analysis on isolated breathing pauses, meaning that any pauses that had another pause in temporal proximity were discarded from the analysis (here, defined as any other pause of at least 5 seconds within -60 to 90 seconds relative to the pause). This meant that we retained 5,910 isolated pauses. Taken together, we included 5,910 pauses in breathing between 5 and 15 seconds from 118 recordings and 848 apnoeas from 64 recordings for analysis of EEG changes.

### 2.5 Time-frequency EEG amplitude

We next computed the time-frequency EEG amplitude for each of the breathing pauses (see Figure 1B). To do so, we first pre-processed the EEG time series. Recording-specific EEG time series were first band-pass filtered between 0.1 and 30 Hz (bidirectional second-order Butterworth filter). We limit our spectral analysis to 30 Hz as infant EEG does typically not encompass higher frequencies such as the gamma band (>30 Hz) (André et al., 2010). EEG time series were then epoched from -90 to 150 seconds around the apnoea and from -90 to 90 seconds around the 5-15-seconds breathing pause onset. We chose 150 and 90 seconds after the pause onset because of the different pause durations; this allowed us to have 60 seconds of data after the pause ended. We next inspected the data quality of these EEG epochs, which was done visually for apnoeas and automated for the 5-15 seconds pauses. Apnoea-related EEG epochs were visually inspected for artefacts including gross movement artefact and electrical interference and were rejected on a channel-wise basis (561 ± 37 epochs were retained). For the 5-15 seconds pauses, we rejected epochs with excessive amplitudes, here defined as values below -500 or higher than 500 µV (4,156 ± 393 epochs were retained).

For each epoch, we computed the time-frequency EEG amplitude. To do this, EEG data were iteratively band-pass filtered at centre frequencies of 1.5 to 29.5 Hz with steps of 1 Hz (i.e., 1.5, 2.5, 3.5, …, 29.5 Hz) and cut-off frequencies at -1 and 1 Hz around the centre frequencies (second-order bidirectional Butterworth filter). Band-pass filtered EEG time series were transformed to the analytic signal using the Hilbert transform, which contains amplitude and phase information in the form of a complex-valued signal. We took the modulus of this analytic signal to retain the amplitude. Time-frequency-amplitude representations were logarithmically transformed before averaging aiming to bring the distribution closer to being Gaussian (Halliday et al., 1995).

### 2.6 Normal breathing

We aimed to test whether the time-frequency EEG amplitude was different between breathing pauses and periods of normal breathing. To define the latter, we searched for periods of normal breathing in the 90 seconds leading up to the breathing pause. We therefore determined the maximal inter-breath interval in time windows of either 30 seconds (for comparison with apnoeas) or 10 seconds (for comparison with 5-15 seconds pauses), shifted the window by 1 second, and computed the maximal inter-breath interval again. This was done up to 15 and 5 sec (apnoea and shorter pauses respectively) before the start of the breathing pause. We then selected the time window in which the maximal inter-breath interval was lowest as normal breathing. Put differently, this was the time windows which had the shortest “pause” between breaths over the entire window, which was 1.16 ± 0.36 seconds (median ± interquartile range over all epochs). The selection of the normal breathing time window was done for every breathing pause individually. Time windows were set to 30 and 10 seconds as time-frequency EEG amplitudes were segmented between -15 and 15 seconds (apnoeas) and -5 and 5 seconds (breathing pauses) at the start/end during the statistical comparisons (see below).

### 2.7 Defining the heart rate and peripheral oxygen saturation outcomes

Before computing the heart rate and oxygen saturation outcomes, we first pre-processed these physiological data. Heart rate values below 40 or above 230 beats per minute and oxygen saturation values above 100% were marked and discarded from the time series. Oxygen saturation was automatically capped at a minimum value of 60% by the recording software.

Heart rate and oxygen saturation changes were defined as the change during the breathing pause relative to the time window used for normal breathing. During the breathing pause, we selected the minimal value of the heart rate and oxygen saturation in the -5 to 60 seconds window relative to the breathing pause end. This longer time window was chosen as the grand average responses showed changes in this time window (Figure 5). EEG and respiratory data of recordings without heart rate and oxygen saturation were not included in this analysis (removing 13 apnoeas and 648 shorter pauses from this analysis).

### 2.8 Statistical analysis

We tested if the time-frequency EEG amplitudes significantly differed between normal breathing and breathing pauses (encompassing either apnoeas or 5-15 seconds pauses). We focused on the start and end of the pause (pause durations are different so these two time-locking procedures may yield different results). Time-frequency EEG amplitudes were segmented between -15 and 15 seconds (apnoeas) and -5 and 5 seconds (breathing pauses) at the start and end. Segments of breathing pauses and normal breathing were statistically compared using sample-wise within-subject *t*-tests with α-levels including false discovery rates of 0.01. We investigated whether time-frequency EEG amplitudes changes were associated with heart rate changes, oxygen saturation changes, breathing pause duration, and post-menstrual age (PMA; sum of gestational age at birth and postnatal age). To do this, we created four linear mixed effects models with EEG amplitude changes as response variable, one for each fixed factor. Infant was included as a random factor on both intercept and slope. Time-frequency EEG amplitude changes were calculated in the time windows of -5 to 5 seconds relative to the breathing pause end, which is where the maximal effect occurred (see Figures 3-5). Time-frequency EEG amplitude changes were averaged over all time and frequency samples for each apnoea/pause. For PMA, we averaged the EEG amplitude changes for every infant (after having averaged over time and frequency), to exclude that the regression models are dominated by infants with many breathing pauses.

## 3 Results

### 3.1 EEG amplitude decreases during both apnoea and shorter pauses in breathing

A total of 848 apnoeas were detected in 64 recordings with an average duration of 17.68 ± 5.22 [15.04-66.48] seconds (median ± interquartile range [range]; Figure 2A). Recordings contained an average of 4 ± 15 [1-105] apnoeas and infants were aged between 32 and 48 weeks, with most infants older than 36 weeks (Figure 2B). In 118 recordings, we additionally identified 5,910 isolated breathing pauses between 5 to 15 seconds with a median duration of 6.04 ± 1.56 [5.04-14.96] seconds (Figure 2C). We found 47 ± 37 [1-152] shorter breathing pauses per recording and infants were aged between 32 and 48 weeks (Figure 2D).

**Figure 2.**
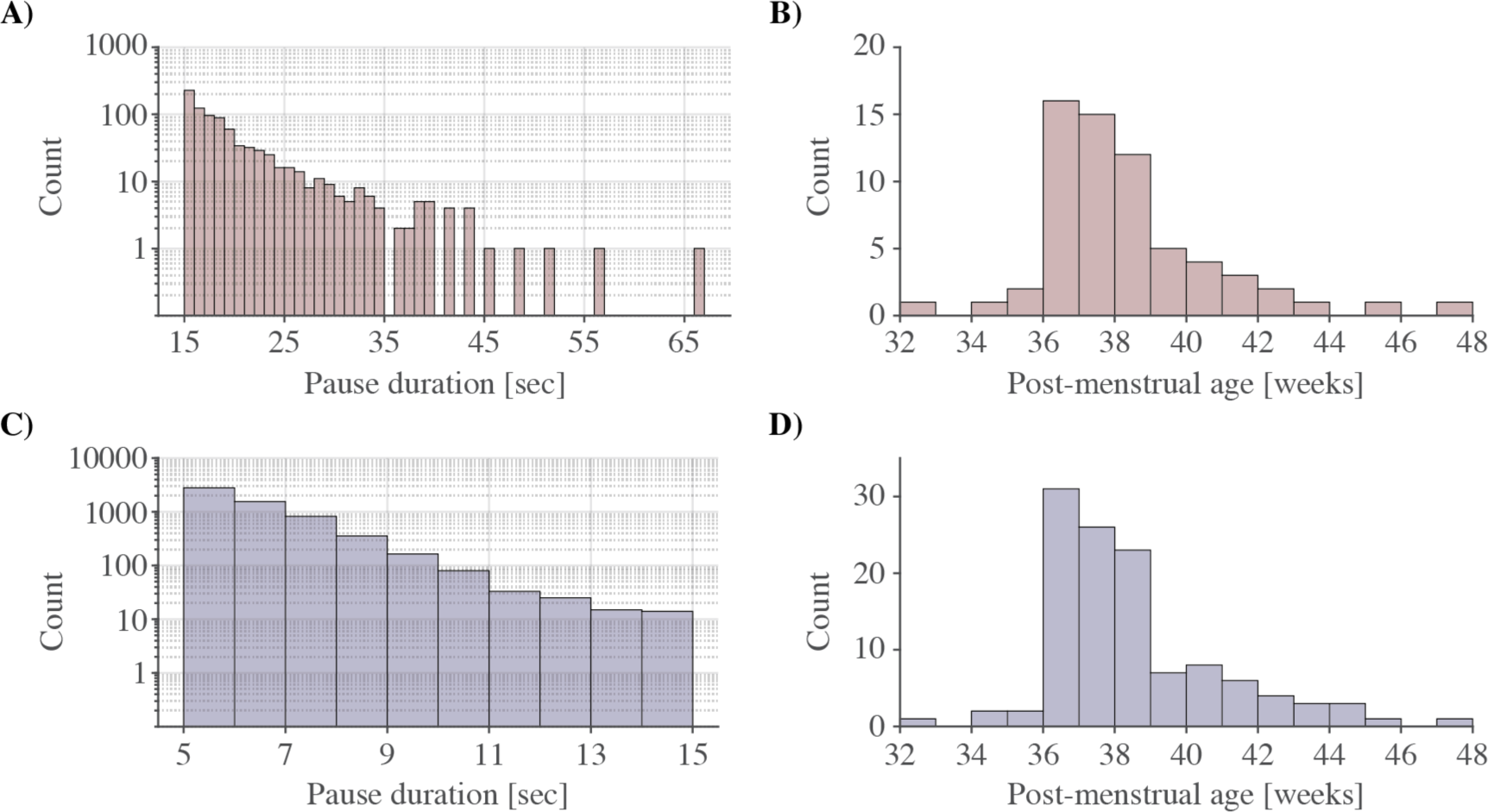
Apnoea/breathing pause durations and post-menstrual ages of infants experiencing cessation of breathing. Number of **A)** apnoeas as a function of the pause duration (shown on a logarithmic scale to improve visualisation) and **B)** post-menstrual ages at time of the recording of infants with apnoeas (n = 64). Number of **C)** breathing pauses between 5 and 15 sec as a function of the pause duration and **D)** post-menstrual ages of infants with these shorter breathing pauses (n = 118).

**Figure 3.**
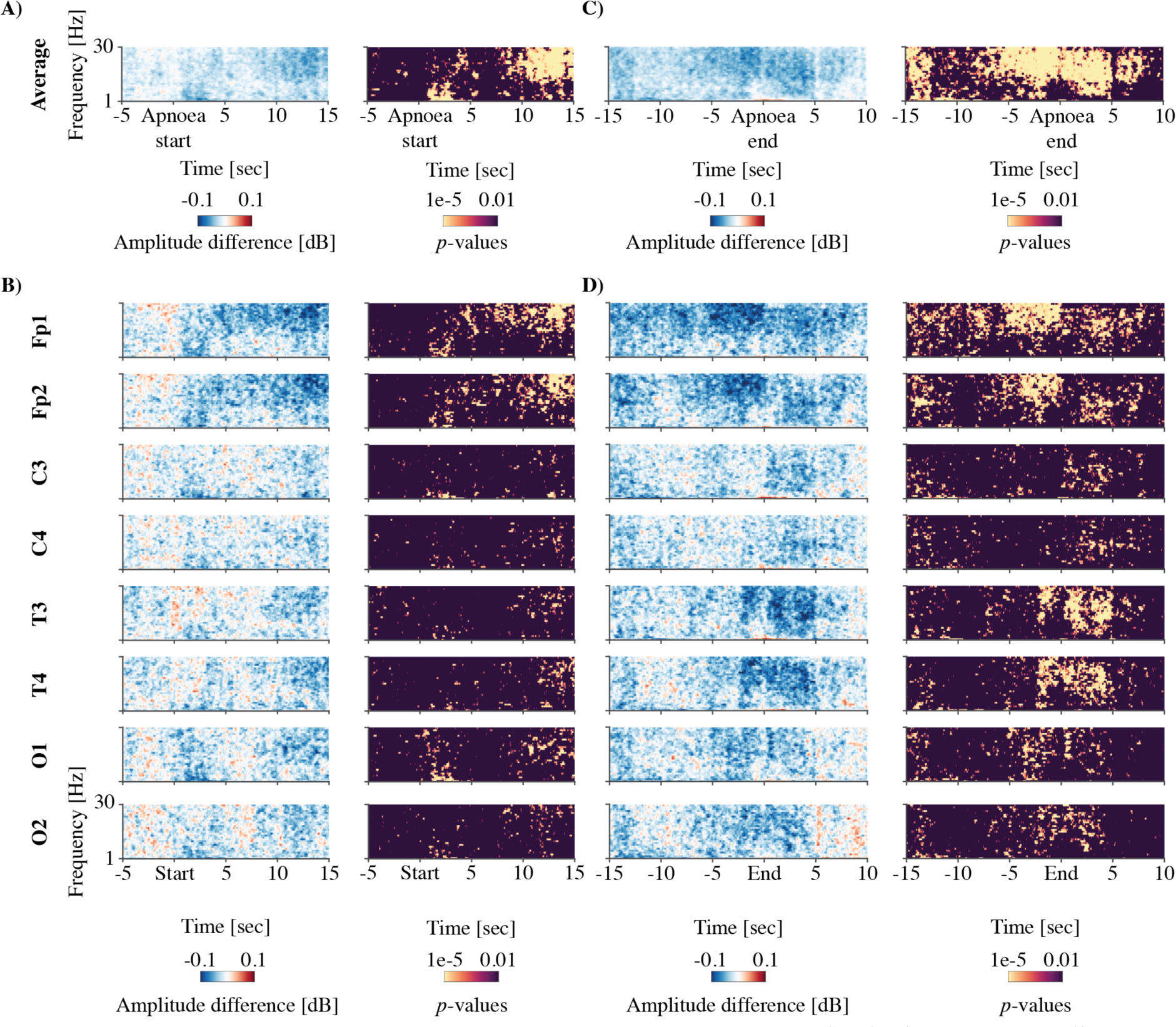
Apnoea-related time-frequency EEG amplitude changes. EEG amplitude changes across all apnoeas of at least 15 seconds in duration (n=848 in 64 recordings). Mean EEG amplitude during the apnoea is compared to normal breathing in the time-frequency plane. Due to the varying apnoea durations we time-locked to both the start of the apnoea **(A-B)** and the end of the apnoea **(C-D)**. **A)** and **C)** show the amplitude change averaged across all channels and **B)** and **D)** are channel specific. Each panel shows the amplitude difference between the apnoea and normal breathing on the left (blue and red plot) and the corresponding statistical difference at each time- frequency point on the right (yellow and black plot). A negative, blue-coloured amplitude difference means a decrease during the apnoea. Statistical significance is set to a false discovery rate of 0.01.

We compared the time-frequency EEG amplitudes during apnoea and breathing pauses between 5 and 15 seconds with periods of normal breathing. Due to varying durations, we assessed EEG amplitude changes time-locked to both the start and end of the apnoeas and shorter pauses in breathing. Immediately after the apnoea started the mean EEG amplitude decreased significantly relative to normal breathing and this suppression continued during the apnoea (Figure 3A). This change in EEG amplitude was particularly marked for frontal channels (Fp1 and Fp2) (Figure 3B). To a lesser extent amplitude changes were also observed at parietal (C3 and C4), temporal (T3 and T4), and occipital channels (O1 and O2) channels. Time-locking to the end of the apnoea, decreases in EEG amplitude were more prominent, occurred across all frequencies, and continued to occur for approximately 5 seconds after the apnoea had ended (Figure 3C). Similar to the start of the apnoea, whilst the decrease in EEG amplitude was most pronounced at frontal electrodes, significant decreases in EEG amplitude were observed across all channels (Figure 3D).

Similarly, for pauses in breathing between 5 and 15 seconds, we found that the average EEG amplitude over all channels decreased during the pause (Figure 4A). Moreover, in the 5 seconds before the pause start were characterised by a decrease at lower frequencies (∼5 to 20 Hz) and increase at higher frequencies around 30 Hz, particularly at temporal channels (Figure 4B). Immediately after the pause started, the time-frequency EEG amplitude reduced for low frequencies (typically below ∼20 Hz) across all channels (Figure 4B). Time-locking to the end of the pause, a significant decrease in EEG amplitude was observed during the pause in breathing but, unlike apnoeas, this decrease in EEG amplitude does not remain after the end of the pause in breathing (Figure 4C-D). In fact, in the average and in some channels, there was a significant increase in EEG amplitude at the end of the pause in breathing.

**Figure 4.**
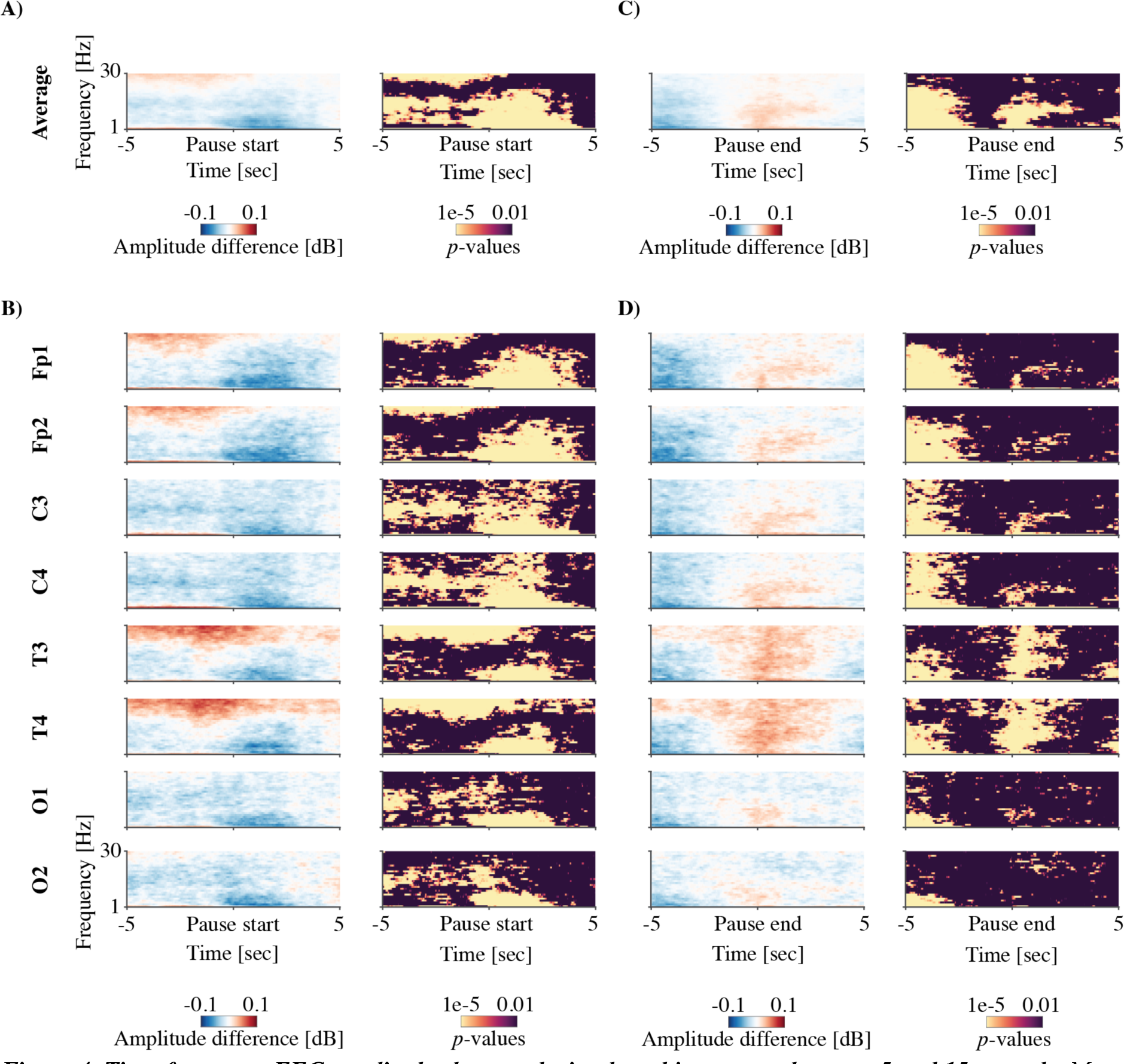
Time-frequency EEG amplitude changes during breathing pauses between 5 and 15 seconds. Mean EEG amplitudes during the pause are compared to normal breathing for 5,910 pauses in breathing in 118 recordings. EEG amplitude changes were assessed time-locked to pause start (**A-B**) and end (**C-D**). **A)** and **C)** show the amplitude change averaged across all channels and **B)** and **D)** are channel specific. Each panel shows the amplitude difference between the pause in breathing and normal breathing on the left (blue and red plot) and the corresponding statistical difference at each time-frequency point on the right (yellow and black plot). A negative (blue) amplitude difference means a decrease during the apnoea. Statistical significance is set to a false discovery rate of 0.01.

**Figure 5.**
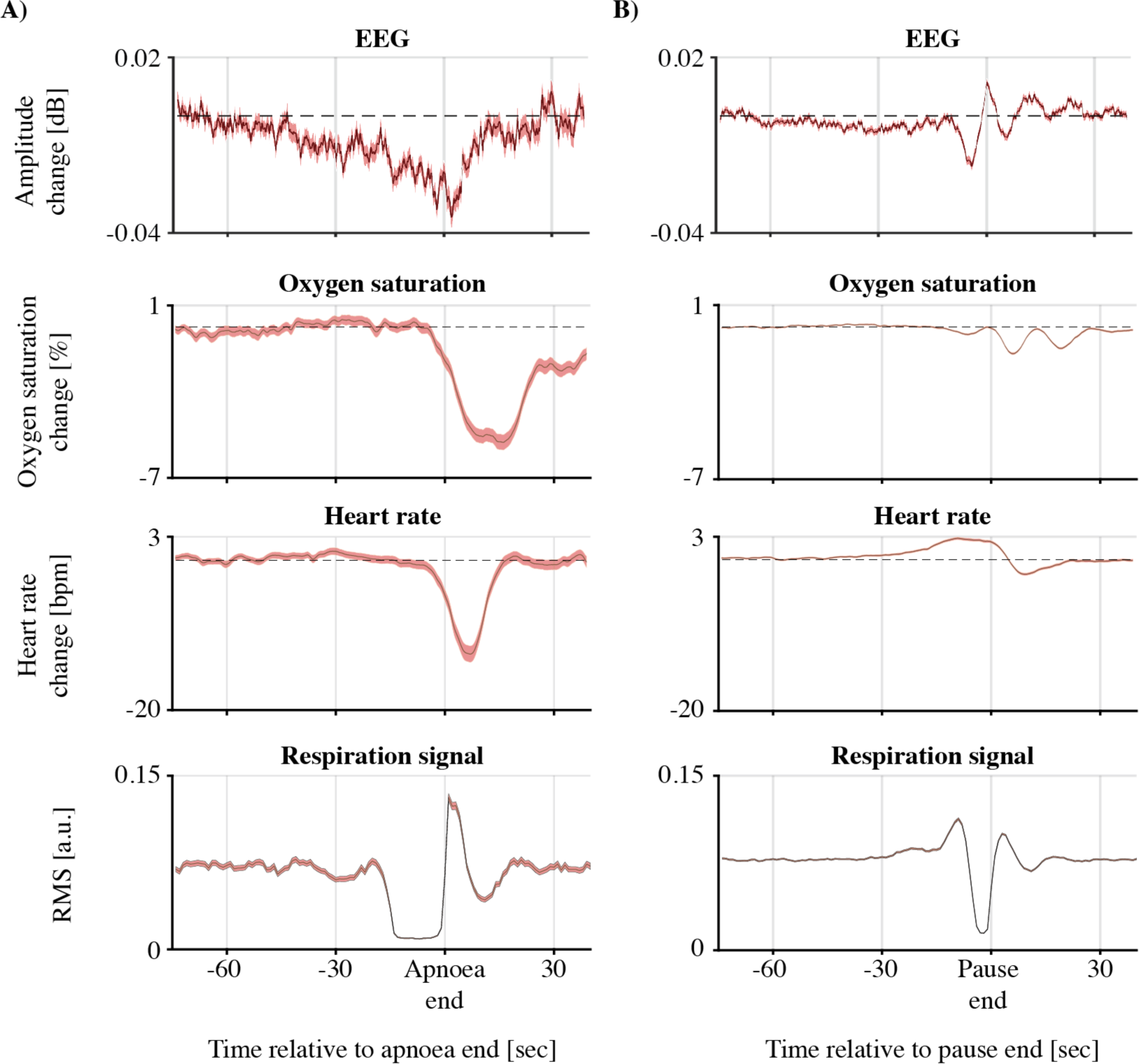
Time-resolved EEG amplitudes and vital signs. Time series were time-locked to the end of the **A)** apnoeas and **B)** breathing pauses between 5 and 15 seconds. Continuous graphs are the mean over all pauses with the shaded surfaces indicating the standard error of the mean. Time-frequency EEG amplitude and vital signs changes are the mean over all pauses (either apnoea [n=848] or 5-15 seconds pauses [n=5,910]) and channels. Time-resolved EEG amplitudes were averaged over frequencies and channels. Respiration signal was defined as a moving root-mean-square (RMS) in one-second windows. The moving average of each breathing pause was normalised to its own norm before averaging over breathing pauses.

### 3.2 Time-resolved EEG amplitude changes are related to changes in heart rate and oxygen saturation

To compare EEG amplitude changes with alterations in heart rate and oxygen saturation, we computed the time-resolved EEG amplitude by pooling the EEG amplitudes over all frequency bands and channels (Figure 5; Figure S1 show the time-resolved EEG amplitude for each channel separately). As expected, for both apnoea and short breathing pauses, on average heart rate and oxygen saturation decreased (Figure 5). Change in EEG amplitude was positively associated with change in heart rate for both apnoea (β*_slope_*= 0.0009; *p* < 0.0001; Figure 6A) and 5-15 seconds breathing pauses (β*_slope_* = 0.0004; *p* = 0.0001; Figure 6B). Change in EEG amplitude was significantly correlated with change in oxygen saturation during apnoeas (β*_slope_* = 0.0015; *p* = 0.0001) but not for shorter breathing pauses (β*_slope_* = -0.0001; *p* = 0.31). Finally, the EEG amplitude changes did not significantly relate to either pause/apnoea duration or age (Figure 6).

**Figure 6.**
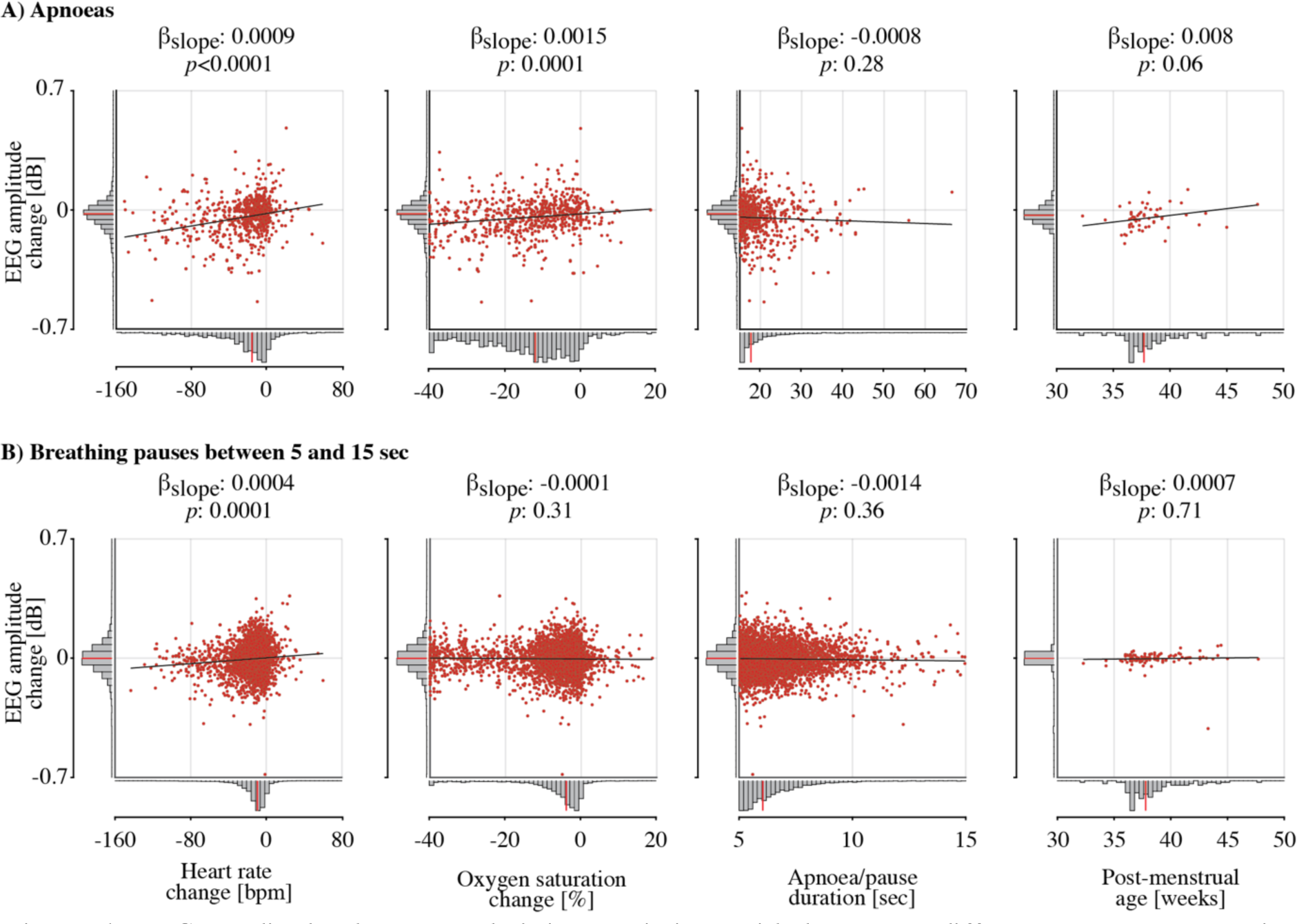
EEG amplitude changes and their associations with heart rate difference, oxygen saturation difference, breathing pause/apnoea duration, and post-menstrual age. EEG amplitude changes for the A) apnoeas and B) breathing pauses between 5 and 15 seconds compared with each of the predictor variables. Heart rate and oxygen saturation changes are computed as the minimal value in the time window of -5 to 60 seconds relative to apnoea and breathing pause end compared with the baseline. Baseline window was defined when the maximal inter-breath interval was lowest in the 90 seconds leading up to the pause. Red data points are the data from a single pause/apnoea in the case of heart rate change, oxygen saturation change and pause duration. For comparison with post-menstrual age, we averaged the amplitude changes over all pauses of a recording. The solid black graphs indicate the model fit. Histograms on the x- and y-axes show the predictor and response variables’ distributions (including their median indicated by the red line).

## 4 Discussion

We investigated how breathing pauses alter cortical activity in infants. Consistent with our hypothesis, EEG amplitude significantly decreased during both apnoea and shorter pauses in breathing. Change in EEG amplitude was significantly associated with change in heart rate and oxygen saturation but not apnoea/pause duration or the age of the infant.

A reduction in cortical activity during apnoea is in line with previous studies (Bridgers et al., 1985; Deuel, 1973; Low et al., 2012; Wulbrand and Bentele, 1994). Whilst most of these studies have reported how cortical activity changes based on relatively small sample sizes (i.e., <10 infants), we demonstrated that these results are consistent in a much larger sample size.

EEG records the post-synaptic activity of cortical neuronal populations, meaning that reduced EEG amplitude implies a decrease in neuronal activity and/or desynchronisation. In rodents, ionic concentrations change significantly during periods of cerebral deoxygenation, where adenosine increases, inhibiting the generation of postsynaptic potentials (Ilie et al., 2006). Adenosine may change the calcium and potassium concentrations, reducing the neuron’s excitability (Luhmann and Kral, 1997; Nabetani et al., 1995). Neural excitability is also linked to tissue acidity (Chesler, 2003). When pH decreases following rising carbon dioxide levels (Chesler, 2003), the concentration of neuromodulator adenosine is increased (Ito et al., 2003). The reduction in cortical activity may have a neuroprotective function during apnoea (Low et al., 2012) as the brain may aim to avoid metabolic failure and irreversible cell damage by conserving energy (Nabetani et al., 1995).

Although the brain may have mechanisms in place to protect itself against metabolic and homeostatic changes, it is possible that the reduced EEG amplitude may still have a negative impact on brain development, particularly if it occurs frequently. Brain activity is essential for brain development, and blocking, reducing or changing the patterning of this activity can result in disruptions to brain development in animal and computational models (Ghosh and Shatz, 1992; Hartley et al., 2020; Schmidt and Eisele, 1985; Tolner et al., 2012). If episodes of apnoea frequently occur and lead to changes in brain activity, this may result in poorer neurodevelopmental outcomes (Janvier et al., 2004; Pillekamp et al., 2007; Poets et al., 2015). Further studies investigating changes in brain development longitudinally compared with apnoea frequency are warranted.

Interestingly, while the changing oxygen and carbon dioxide concentrations may decrease brain activity, we observed changes in cortical activity immediately after the pause started. These may not be the result of neurotransmitter alternations from changes in gas concentrations as these rely on slower dynamics (Figure 5). Alternatively, cortico-respiratory coupling may change and be responsible for the immediate alterations in cortical activity. For example, in rodents coupling between the nasal airstream and olfactory system is established via mechanoreceptors (Ito et al., 2014), where slow respiratory rhythms entrain high-frequency neural oscillations. Coupling is most prominent between breathing and frontal areas (Karalis and Sirota, 2022) which is where we found strongest modulation. Cross-frequency coupling is also evident in adults where the phase of breathing below 1 Hz is related to theta- and beta-band oscillations (Schreiner et al., 2023). This cortical-respiratory coupling is likely to change when breathing is interrupted; however, further work is needed to study this in infants.

Apnoeas in infants often co-occur with bradycardia and oxygen desaturations (Poets and Southall, 1991). In our data, both heart rate and oxygen saturation decreases were apparent following apnoea and EEG amplitude changes were associated with heart rate changes and to a lesser extent with oxygen saturation changes. The correlation with change in heart rate is in contrast to a previous study, where no relationship between EEG frequency changes and heart rate were identified (Curzi-Dascalova et al., 2000). However, this study included only five infants with 77 isolated pauses in breathing and compared heart rate and EEG frequency, rather than amplitude, changes. Bradycardia may be directly caused by the hypoxic effect on the heart during an apnoea, including stimulation of the carotid body chemoreceptors (Girling, 1972; Henderson-Smart et al., 1986). When this bradycardia is severe, it may also elicit a decrease in cardiac output together with lower cerebral blood flow and systemic blood pressure. Blood pressure regulation within the brain is not well developed in infants (Weindling and Kissack, 2001) and hence a decrease in systemic blood pressure may result in cerebral hypo-perfusion. This may contribute to the change in brain activity during the pause in breathing. In our data we cannot dissociate whether the EEG suppression is due to the bradycardia (and decrease in cardiac output/cerebral blood flow) or the combination of apnoea and bradycardia. Interestingly, the rate of cerebral blood flow changes may be region dependent which is shown in animal studies investigating asphyxia (Stonestreet et al., 1982). There, blood flow decreases more rapidly in rostral compared to caudal areas of the central nervous system (Greisen, 1992), which may be why we observed that the EEG amplitude changes were most profound in the frontal cerebral areas.

To a lesser extent than with heart rate, we observed positive associations between EEG amplitude changes and oxygen saturation changes (only for apnoeas and not for pauses between 5 to 15 seconds). However, a limitation of our recording of oxygen saturation is that the minimum value was truncated at 60%, meaning that the changes may have been even larger than the maximal change of 40%. Moreover, oxygen saturation was measured peripherally, and changes in cerebral oxygenation may not reflect those in the periphery (Urlesberger et al., 2010). Studies using near-infrared spectroscopy (NIRS) have found that, during apnoea, changes in peripheral oxygenation are not always correlated with changes in cerebral oxygenation, particularly for less severe events (Watkin et al., 1996), and that in many cases cerebral oxygenation is maintained above 60% (Schmid et al., 2015). This is due to vasodilation occurring following rising carbon dioxide and lowering oxygen levels and highlights a limitation of our work in that we did not record changes in carbon dioxide levels.

Future work should combine EEG and NIRS recordings during apnoea, in addition to measuring blood carbon dioxide levels, to further study this mechanism.

Consistent with a previous study (Fenichel et al., 1980), we did not find a relationship between breathing pause duration and EEG amplitude changes. Although changes in physiology are correlated with the duration of an apnoea, there is wide variation, with some comparatively short apnoeas leading to profound oxygen desaturations and bradycardia (Upton et al., 1991). We also did not find a relationship between EEG amplitude changes and age. Whilst the youngest infant included in the study was 31 weeks PMA, the majority of the infants were studied when they were above 36 weeks. Although not significant, there was a trend towards greater amplitude suppression during apnoea in infants of younger ages and so further investigations in this area including younger premature infants are needed. Another factor that we have not considered is sleep staging, which is important as it is closely related to metabolic demands. For example, most apnoeas happen during REM sleep, which is a state that requires most metabolic energy (Lehtonen and Martin, 2004). Moreover, our analysis did not specifically differentiate the type of apnoea the infants were experiencing: apnoeas can be central: where there is cessation of the drive to breath; obstructive: where there is an obstruction of the airway; or mixed: where there is a combination of the two. Analysis of the shorter pauses in breathing focused on the signal from the thoracic band and so included central or mixed pauses in breathing. Future studies should examine if these different apnoea types have differential effects on brain activity. Finally, we focused our analysis on changes in EEG amplitude during isolated pauses in breathing (with no other short pauses in breathing in the preceding -60 seconds or the 90 seconds after the pause). This enabled us to clearly attribute changes in brain activity to the single pauses and compare with other changes in physiology. However, clusters of pauses in breathing, such as in periodic breathing, can occur frequently in preterm infants, and future work should compare changes in brain activity during isolated pauses with that during clustered pauses.

In summary, we found that cortical activity decreases during breathing pauses over the whole cortex. This modulation of cortical activity was observed for both apnoeas (pauses greater than 15 seconds) and shorter breathing pauses. Although the decrease in cortical activity may not be harmful, it is unknown if this extrapolates to repeated apnoeas which result in chronic intermittent hypoxia, and how this affects long-term brain development. Moreover, our study mostly focused on term-corrected infants. Due to the rapid functional and structural changes in the preterm period, neurophysiological effects may be larger in preterm infants and further investigation may shed light on critical developmental windows for the effects of apnoea on brain development.

## Data and code availability

The data that support the study findings are available upon reasonable request. Data sharing requests should be directed to. Requests for code sharing should be directed to.

## Author contributions

**Coen S. Zandvoort:** Conceptualization, Methodology, Software, Formal analysis, Writing – Original Draft, Writing – Review & Editing, Visualization; **Anneleen Dereymaeker:** Investigation, Data Curation, Writing – Review & Editing; **Luke Baxter:** Methodology, Writing – Review & Editing; **Katrien Jansen:** Investigation, Data Curation, Writing – Review & Editing; **Gunnar Naulaers:** Investigation, Writing – Review & Editing; **Maarten de Vos:** Methodology, Writing – Review & Editing; **Caroline Hartley:** Conceptualization, Methodology, Writing – Original Draft, Writing – Review & Editing, Supervision, Funding acquisition

## Acknowledgements and funding

The authors would like to thank all parents and infants involved in the studies and staff at the UZ Leuven NICU in Leuven who helped with data collection. We would also like to thank Ayesha Ali for assistance with the classification labelling in an early form of the logistic regression model used to identify short pauses in breathing. This study is funded by a Wellcome Trust/Royal Society Sir Henry Dale Fellowship (grant number: 213486/Z/18/Z), awarded to CH.

## Declaration of competing interests

The authors declare no conflicts of interest.

## Notes

### Competing Interest Statement

The authors have declared no competing interest.

## References

1. Adjei, T., Purdy, R., Jorge, J., Adams, E., Buckle, M., Fry, R.E., Green, G., Patel, C., Rogers, R., Slater, R., 2021. New method to measure interbreath intervals in infants for the assessment of apnoea and respiration. BMJ Open Respiratory Research 8, e001042.

2. André, M., Lamblin, M.-D., d’Allest, A.-M., Curzi-Dascalova, L., Moussalli-Salefranque, F., Tich, S.N.T., Vecchierini-Blineau, M.-F., Wallois, F., Walls-Esquivel, E., Plouin, P., 2010. Electroencephalography in premature and full-term infants. Developmental features and glossary. Neurophysiologie Clinique/Clinical neurophysiology 40, 59–124.

3. Blackmon, L.R., Batton, D.G., Bell, E.F., Engle, W.A., 2003. Apnea, sudden infant death syndrome, and home monitoring. Pediatrics 111, 914–914.

4. Bridgers, S.L., Ment, L.R., Ebersole, J.S., Ehrenkranz, R.A., 1985. Cassette electroencephalographic recording of neonates with apneic episodes. Pediatric Neurology 1, 219–222.

5. Chesler, M., 2003. Regulation and modulation of ph in the brain. Physiological Reviews 83, 1183–1221.

6. Choi, S.H., Lee, J., Nam, S.K., Jun, Y.H., 2021. Cerebral oxygenation during apnea in preterm infants: Effects of accompanying peripheral oxygen desaturation. Neonatal Medicine 28, 14–21.

7. Curzi-Dascalova, L., Bloch, J., Vecchierini, M.-F., Bedu, A., Vignolo, P., 2000. Physiological parameters evaluation following apnea in healthy premature infants. Neonatology 77, 203–211.

8. Deuel, R.K., 1973. Polygraphic monitoring of apneic spells. Archives of Neurology 28, 71–76.

9. Eichenwald, E.C., Aina, A., Stark, A.R., 1997. Apnea frequently persists beyond term gestation in infants delivered at 24 to 28 weeks. Pediatrics 100, 354–359.

10. Elder, D.E., Campbell, A.J., Galletly, D., 2013. Current definitions for neonatal apnoea: Are they evidence based? Journal of Paediatrics and Child Health 49, E388–E396.

11. Fenichel, G.M., Olson, B.J., Fitzpatrick, J.E., 1980. Heart rate changes in convulsive and nonconvulsive neonatal apnea. Annals of Neurology: Official Journal of the American Neurological Association and the Child Neurology Society 7, 577–582.

12. Finer, N.N., Higgins, R., Kattwinkel, J., Martin, R.J., 2006. Summary proceedings from the apnea-of-prematurity group. Pediatrics 117, S47-S51.

13. Ghosh, A., Shatz, C.J., 1992. Involvement of subplate neurons in the formation of ocular dominance columns. Science 255, 1441–1443.

14. Girling, D.J., 1972. Changes in heart rate, blood pressure, and pulse pressure during apnoeic attacks in newborn babies. Archives of Disease in Childhood 47, 405–410.

15. Greisen, G., 1992. Effect of cerebral blood flow and cerebrovascular autoregulation on the distribution, type and extent of cerebral injury. Brain Pathology 2, 223–228.

16. Halliday, D., Rosenberg, J., Amjad, A., Breeze, P., Conway, B., Farmer, S., 1995. A framework for the analysis of mixed time series/point process data-theory and application to the study of physiological tremor, single motor unit discharges and electromyograms. Progress in Biophysics and Molecular Biology 64, 237.

17. Hartley, C., Farmer, S., Berthouze, L., 2020. Temporal ordering of input modulates connectivity formation in a developmental neuronal network model of the cortex. PLoS One 15, e0226772.

18. Henderson-Smart, D., Butcher-Puech, M., Edwards, D.A., 1986. Incidence and mechanism of bradycardia during apnoea in preterm infants. Archives of Disease in Childhood 61, 227–232.

19. Horne, R.S., Sun, S., Yiallourou, S.R., Fyfe, K.L., Odoi, A., Wong, F.Y., 2018. Comparison of the longitudinal effects of persistent periodic breathing and apnoea on cerebral oxygenation in term-and preterm-born infants. The Journal of Physiology 596, 6021–6031.

20. Ilie, A., Ciocan, D., Zagrean, A.-M., Nita, D.A., Zagrean, L., Moldovan, M., 2006. Endogenous activation of adenosine a1 receptors accelerates ischemic suppression of spontaneous electrocortical activity. Journal of Neurophysiology 96, 2809–2814.

21. Ito, H., Kanno, I., Ibaraki, M., Hatazawa, J., Miura, S., 2003. Changes in human cerebral blood flow and cerebral blood volume during hypercapnia and hypocapnia measured by positron emission tomography. Journal of Cerebral Blood Flow & Metabolism 23, 665–670.

22. Ito, J., Roy, S., Liu, Y., Cao, Y., Fletcher, M., Lu, L., Boughter, J., Grün, S., Heck, D., 2014. Whisker barrel cortex delta oscillations and gamma power in the awake mouse are linked to respiration. Nature Communications 5, 3572.

23. Janvier, A., Khairy, M., Kokkotis, A., Cormier, C., Messmer, D., Barrington, K.J., 2004. Apnea is associated with neurodevelopmental impairment in very low birth weight infants. Journal of Perinatology 24, 763–768.

24. Karalis, N., Sirota, A., 2022. Breathing coordinates cortico-hippocampal dynamics in mice during offline states. Nature Communications 13, 467.

25. Lehtonen, L., Martin, R.J., 2004. Ontogeny of sleep and awake states in relation to breathing in preterm infants. Seminars in Neonatology. Elsevier, pp. 229–238.

26. Low, E., Dempsey, E.M., Ryan, C.A., Rennie, J.M., Boylan, G.B., 2012. Eeg suppression associated with apneic episodes in a neonate. Case Reports in Neurological Medicine 2012.

27. Luhmann, H.J., Kral, T., 1997. Hypoxia-induced dysfunction in developing rat neocortex. Journal of Neurophysiology 78, 1212–1221.

28. Martin, R.J., Abu-Shaweesh, J.M., Baird, T.M., 2004. Apnoea of prematurity. Paediatric Respiratory Reviews 5, S377–S382.

29. Nabetani, M., Okada, Y., Kawai, S., Nakamura, H., 1995. Neural activity and the levels of high energy phosphates during deprivation of oxygen and/or glucose in hippocampal slices of immature and adult rats. International Journal of Developmental Neuroscience 13, 3–12.

30. Pillay, K., Dereymaeker, A., Jansen, K., Naulaers, G., De Vos, M., 2020. Applying a data-driven approach to quantify eeg maturational deviations in preterms with normal and abnormal neurodevelopmental outcomes. Scientific Reports 10, 1–14.

31. Pillay, K., Dereymaeker, A., Jansen, K., Naulaers, G., Van Huffel, S., De Vos, M., 2018. Automated eeg sleep staging in the term-age baby using a generative modelling approach. Journal of Neural Engineering 15, 036004.

32. Pillekamp, F., Hermann, C., Keller, T., Von Gontard, A., Kribs, A., Roth, B., 2007. Factors influencing apnea and bradycardia of prematurity–implications for neurodevelopment. Neonatology 91, 155–161.

33. Poets, C.F., Roberts, R.S., Schmidt, B., Whyte, R.K., Asztalos, E.V., Bader, D., Bairam, A., Moddemann, D., Peliowski, A., Rabi, Y., Solimano, A., Nelson, H., Investigators, C.O.T., 2015. Association between intermittent hypoxemia or bradycardia and late death or disability in extremely preterm infants. JAMA 314, 595–603.

34. Poets, C.F., Southall, D.P., 1991. Patterns of oxygenation during periodic breathing in preterm infants. Early Human Development 26, 1–12.

35. Sanocka, U., Donnelly, D., Haddad, G., 1988. Cardiovascular and neurophysiologic changes during graded duration of apnea in piglets. Pediatric Research 23, 402–407.

36. Schmid, M.B., Hopfner, R.J., Lenhof, S., Hummler, H.D., Fuchs, H., 2015. Cerebral oxygenation during intermittent hypoxemia and bradycardia in preterm infants. Neonatology 107, 137–146.

37. Schmidt, J., Eisele, L., 1985. Stroboscopic illumination and dark rearing block the sharpening of the regenerated retinotectal map in goldfish. Neuroscience 14, 535–546.

38. Schreiner, T., Petzka, M., Staudigl, T., Staresina, B.P., 2023. Respiration modulates sleep oscillations and memory reactivation in humans. Nature Communications 14, 8351.

39. Stonestreet, B.S., Laptook, A., Schanler, R., William, O., 1982. Hemodynamic responses to asphyxia in spontaneously breathing newborn term and premature lambs. Early Human Development 7, 81–97.

40. Tolner, E.A., Sheikh, A., Yukin, A.Y., Kaila, K., Kanold, P.O., 2012. Subplate neurons promote spindle bursts and thalamocortical patterning in the neonatal rat somatosensory cortex. Journal of Neuroscience 32, 692–702.

41. Upton, C., Milner, A., Stokes, G., 1991. Apnoea, bradycardia, and oxygen saturation in preterm infants. Archives of Disease in Childhood 66, 381–385.

42. Urlesberger, B., Grossauer, K., Pocivalnik, M., Avian, A., Müller, W., Pichler, G., 2010. Regional oxygen saturation of the brain and peripheral tissue during birth transition of term infants. The Journal of Pediatrics 157, 740–744.

43. Usman, F., Marchant, S., Baxter, L., Salihu, H.M., Aliyu, M.H., Adams, E., Hartley, C., 2023. The effect of acute respiratory events and respiratory stimulants on eeg-recorded brain activity in neonates: A systematic review. Clinical Neurophysiology Practice.

44. Watkin, S., Spencer, S., Pryce, A., Southall, D., 1996. Temporal relationship between pauses in nasal airflow and desaturation in preterm infants. Pediatric Pulmonology 21, 171–175.

45. Weindling, C.M., Kissack, M.A., 2001. Blood pressure and tissue oxygenation in the newborn baby at risk of brain damage. Biology of the Neonate 79, 241–245.

46. WHO, 2023. 152 million babies born preterm in the last decade. https://www.who.int/news/item/09-05-2023-152-million-babies-born-preterm-in-the-last-decade. Accessed on 21/12/2023.

47. Williamson, M., Poorun, R., Hartley, C., 2021. Apnoea of prematurity and neurodevelopmental outcomes: Current understanding and future prospects for research. Frontiers in Pediatrics 9, 755677.

48. Wulbrand, H., Bentele, K.H., 1994. Eeg suppression following augmented breaths during apnea. Pediatric Research 36, 62A–62A.

