## Supplementary figures for "Apnoea suppresses brain activity in infants"

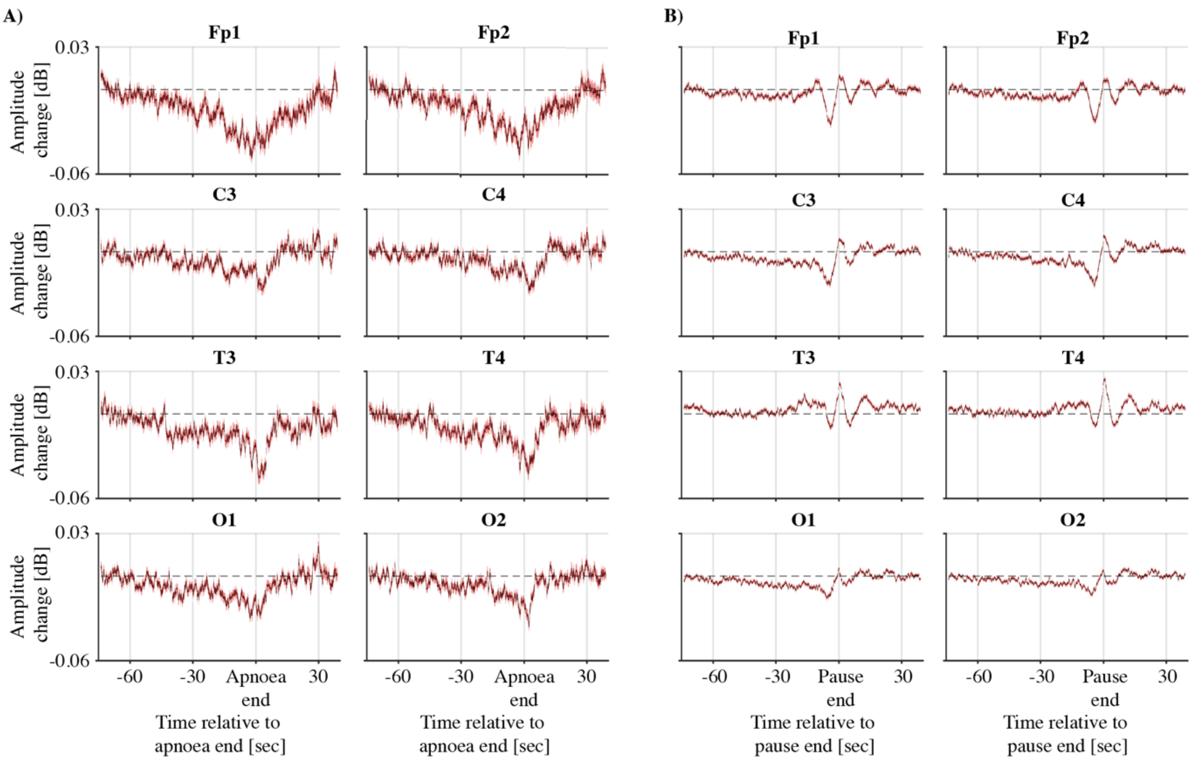


***Figure S1. Time-resolved EEG amplitudes during A) apnoea and B) breathing pauses between 5 and 15 seconds.*** *Amplitudes are pooled over frequencies from 1 and 30 Hz, and time-locked to the end of the apnoea/breathing pause. Continuous thick graphs present the mean amplitude. Shaded surfaces are standard error of the mean over apnoeas/breathing pauses.*
